# Accumulative deamidation of human lens protein γS-crystallin leads to partially unfolded intermediates with enhanced aggregation propensity

**DOI:** 10.1101/2020.02.21.960237

**Authors:** Calvin J. Vetter, David C. Thorn, Samuel G. Wheeler, Charlie Mundorff, Kate Halverson, John A. Carver, Larry L. David, Kirsten J. Lampi

## Abstract

Age-related cataract is a major cause of blindness worldwide. Yet, the molecular mechanisms whereby large, light scattering aggregates form is poorly understood, because of the complexity of the aggregates isolated from human lenses. The predominant proteins in the lens are structural proteins called crystallins. The γS-crystallin is heavily modified in cataractous lenses by deamidation, which introduces a negative charge at labile asparagine residues. The effects of deamidation at asparagines, N14, N76, and N143, were mimicked by replacing the asparagine with aspartate using site-directed mutagenesis. The effects of these surface deamidations on the stability, unfolding, and aggregation properties of γS were determined using dynamic light scattering, chemical and thermal-denaturation, and hydrogen-deuterium exchange with mass spectrometry. We found that a small population of all the deamidation mimics aggregated directly into large light scattering bodies with a radius greater than 10 nm that contributed 14-60% of the total scattering intensity compared to 7% for WT under the same conditions. A possible mechanism was identified under partially denaturing conditions, where deamidation led to significantly more rapid unfolding and aggregation particularly for N76D compared to WT. The triple mutant was further destabilized, reflecting the enhanced aggregation properties of N14D and N143D. Thus, the effects of deamidation were both site-specific and cumulative. αA-crystallin was ineffective at acting as a chaperone to prevent the aggregation of destabilized, deamidated γS. It is concluded that surface deamidations, while causing minimal structural disruption individually, progressively destabilize crystallin proteins, leading to their unfolding and precipitation in aged and cataractous lenses.

## Introduction

In mammals, the eye lens plays a crucial role in vision via transmitting, refracting and focusing light onto the retina. Lens functionality is maintained via a high concentration of crystallin proteins (up to 300-500 mg/mL in the center of the lens) that are arranged in a supramolecular array with short-range order (1). The lens is a unique organ in that it has no blood supply and there is no protein turnover in its fiber cells because they lack the organelles for protein synthesis and degradation. As a result, crystallins are long-lived proteins (2).

Cataract arises from opacification and concomitant light scattering associated with the unfolding, aggregation and precipitation of crystallins. Cataract is the major cause of blindness in the world and is the leading cause of low vision in the United States (3). Age-related cataract is the most prevalent type although many crystallin mutations give rise to congenital and early-onset cataract. The molecular mechanism(s) whereby cumulative modifications of crystallins lead to the formation of large, light scattering aggregates in the aging lens is poorly understood, because of the complexity of the insoluble aggregates isolated from human lenses (4–9).

There are three types of mammalian crystallins, α, β and γ, which are all compromised of a variety of isoforms (10, 11). The two α-crystallins (A and B) are small heat-shock molecular chaperone proteins. The β- and γ-crystallins are homologous to each other and are not related to the α-crystallins. The β- and γ-crystallins are β-sheet-rich, two-domain proteins with the β-sheets adopting a Greek key arrangement. The Greek key motif contains four β-strands. In γS-crystallin (γS), each domain comprises two Greek key motifs, i.e. a total of eight β-strands (Fig. 1 – the NMR structure of γS). The domains in the γ-crystallins are monomers with their domains linked by a connecting peptide that is bent, whereas in β-crystallins the connecting peptide is either bent or extended leading to oligomer formation with hydrophobic interactions stabilizing the interface between the domains (12–15).

**Figure 1:**
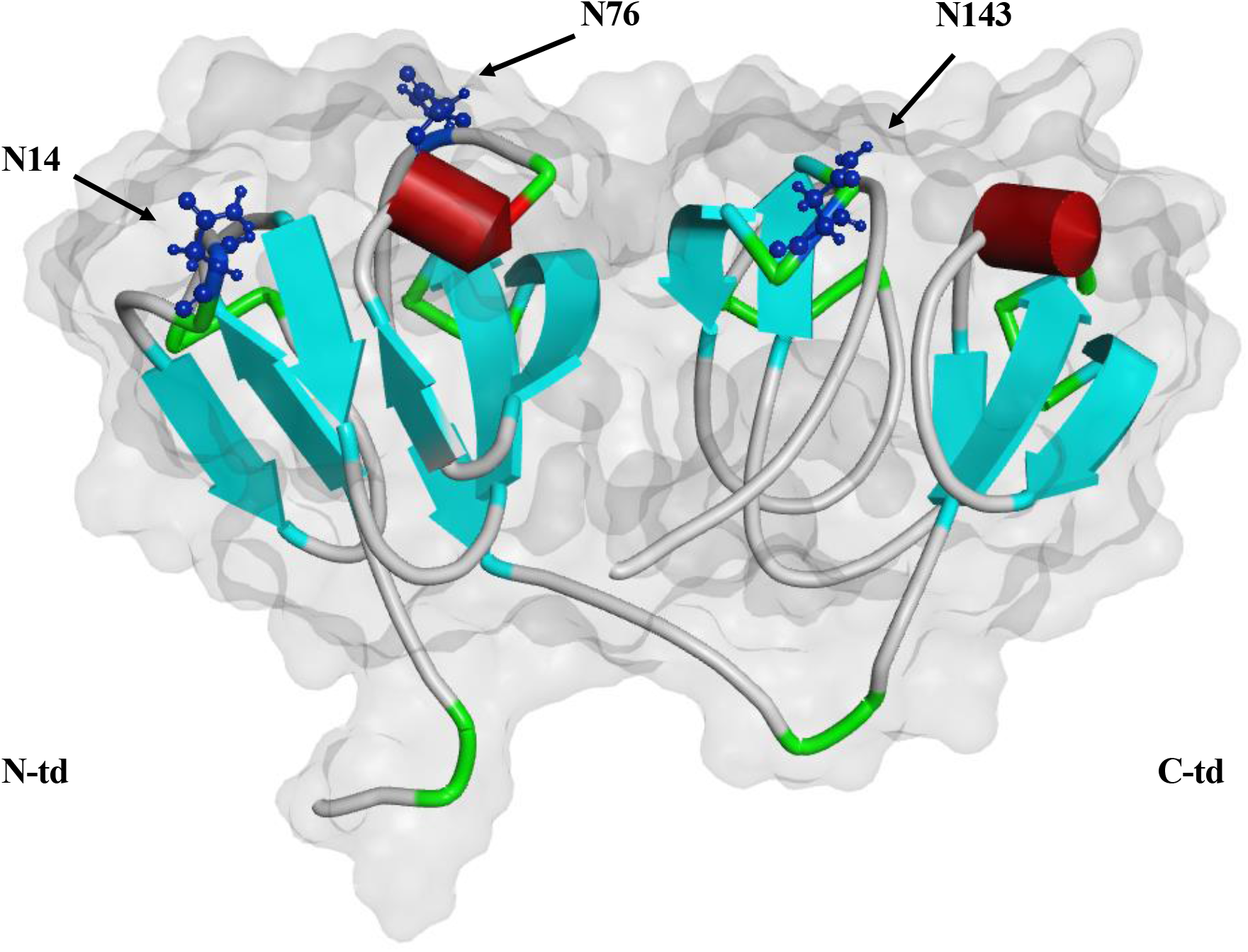
Location of Asn residues 14, 76, and 143 in human γS. In order to mimic deamidation, site-directed mutagenesis was used to replace Asn with Asp at residues 14, 76, and 143. Asparagine residues 14 and 76 are located on surface of the N-terminal domain (N-td) and residue 143 is located on the surface of the C-terminal domain (C-td). Residues of interest are shown in ball and stick structure. Grey area represents solvent accessibility with a probe radius of 1.4, cyan arrows represent β-sheets, red cylinders represent α-helices and green represents coil and turn structures as generated with Discovery Studio Visualizer (BIOVIA, San Diego, CA). (PDB: 2M3T)

Due to the absence of protein turnover, the lens is an incubator that accumulates multiple crystallin modifications over the individual’s lifetime. Deamidation is a major age-related crystallin modification associated with insolubilization of crystallins *in vivo* (6, 16, 17). Deamidation sites at buried interfaces between β- and γ-crystallin domains, while disruptive to protein stability, are not the most prevalent deamidation sites found *in vivo.* The most prevalent are in solvent-exposed regions (6, 18, 19). *In vivo,* γS undergoes significant deamidation with age, in this case the conversion of asparagine residues, principally at surface residues (N14, N76 and N143), each to four possible isomers of aspartic acid, i.e. L-Asp, L-isoAsp, D-Asp and D-isoAsp (20–22). The location of these deamidation sites are indicated on the solution NMR-derived structure of γS (Fig. 1) (23). At these labile surface Asn residues, levels range from 25-60% at N14, 50-80% at N76, and 20-90% at N143 in γS implying a strong association between these deamidation sites and cataract formation (6, 18, 19, 22, 24, 25). The extent of deamidation in the entire crystallin mixture, including these three γS sites (particularly N76 and N143), is significantly greater in cataract lenses compared to age-matched, non-cataract controls, and significantly greater in the insoluble proteins than the soluble proteins from these lenses, implying that these modifications contribute to cataract formation (6, 18, 22, 24, 25). It is expected that the other isomers would affect the protein structure even more.

Previous studies have investigated the effects of deamidation of γS by mimicking deamidated residues with L-Asp, replacing L-Asn via mutagenesis and introducing a negative charge at the site of modification (7, 26). We recently reported that single-point deamidations on the surface of γS can alter its conformational dynamics, providing a potential mechanism for the unfolding and insolubilization of deamidated crystallins in the eye lens (27). In this present study, the effects of deamidation (conversion to L-Asp) at each these three sites, along with the triple mutant (N14D/N76D/N143D, TM), was investigated to determine the accumulative effects of the surface deamidations on the stability, unfolding, and aggregation properties of γS. We found that a small population of all the deamidation mimics aggregated into large light scattering bodies. A possible mechanism was identified under partially denaturing conditions, where deamidation led to more rapid unfolding and aggregation particularly for N76D compared to WT. The triple mutant was further destabilized, reflecting the enhanced aggregation properties of N14D and N143D. Thus, the effects of deamidation were both site-specific and cumulative. Furthermore, αA-crystallin was ineffective at acting as a chaperone to prevent the aggregation of destabilized, deamidated γS. It is concluded that surface deamidations, while causing minimal structural disruption individually, progressively destabilize crystallin proteins, leading to their unfolding and precipitation in aged and cataractous lenses.

## Experimental

### Recombinant expression and purification of human γS and deamidation mimics

The plasmid containing gene for γS wild-type was obtained from the plasmid repository at DNASU (https://dnasu.org/DNASU/Home.do). Single deamidation mimics of N14, N76, N143, plus the triple mimic of all three sites were generated via site-directed mutagenesis using the QuikChangeXL Kit (Aglient Technologies, CA). Protein was expressed in a pE-SUMO (Small Ubiquitin-like Modifier) vector containing a N-terminal 6XHis tagged fusion protein (LifeSensors Inc., PA) using *Escherichia coli.* Culture of cells took place at 37 °C, protein expression was induced with IPTG and growth continued at 37 °C for 4 h. Purification of the protein used HisPur Cobalt Resin following the manufacturer’s protocol (Thermo Fisher Scientific, Waltham, MA.). The removal of the His Tag and SUMO was performed using Ulp-1 protease (Ubl-specific protease 1). Protein was then run through HisPur Cobalt Resin again to remove the tag. The protein was aliquoted, 2.4 mM dithiothreitol (DTT) was added and samples stored at −80 °C. Typical yields of protein were several mg per 100 mL of *E. coli* culture. The purity and masses of all expressed proteins were confirmed by mass spectrometry and gel electrophoresis to match predicted masses and to be greater than 95% pure. Protein concentration was determined by measuring absorbance at 280 nm and calculating concentrations using the protein extinction coefficient calculated using the ExPasy ProtParam tool (human γS 42,860 cm^-1^ M^-1^).

### Dynamic light scattering (DLS) of WT and deamidated γS

Proteins were concentrated to 1 or 5 mg/mL using centrifugation (Amicon Ultra-15 Filter Units (10 kDa MWCO), Millipore Sigma, Burlington, MA). Samples were exchanged into Buffer A (29 mM Na_2_HPO_4_, 29 mM NaH_2_PO_4_, 100 mM KCl, 1 mM EDTA and 1 mM DTT) using dialysis and cleared of large particulate matter using 0.1 μm pore size filters (Ultrafree-MC Centrifugal Filter Units, Millipore Sigma). Samples of 30 uL were loaded into a 384 well plate in replicates of 5 for each γS and the DLS measured at 25 °C using a Dynapro Plate Reader III (Wyatt Technology Corp., Santa Barbara, CA). The time-dependent fluctuations of scattered light intensity were measured and the translational diffusion coefficient (D_t_) was calculated by the autocorrelation function of the DYNAMICS software (Wyatt Tech.). The D_t_ was used to calculate the hydrodynamic radii (*R_h_*) (20). The percent of DLS intensity was plotted for the distributions of different size species in an intensity-weighted regulation graph. Sizes were distributed into different peaks of 1-10 nm, 10-100 nm, and 100-1000 nm and the percent polydispersity (%Pd) was calculated as the distribution of radii within each peak. Weight average molar masses estimated from DLS (M_w_-R) were predicted from the *R*_h_ and assuming a sphere by DYNAMICS software.

### Size-exclusion chromatography and multi-angle light scattering (SEC-MALS) of WT and deamidated γS

WT and deamidated γS were buffer exchanged and concentrated as described for DLS above. The *Mw* were determined from the measured Rayleigh light scattering (Dawn Heleos II and ASTRA VII Software, Wyatt Tech.). A dn/dc of 0.193 and absorbance at 280 nm or a refractive index detector were used to determine concentration (Optilab REX, Wyatt Tech.) (7). Fractions eluting under each peak were analyzed by SDS-PAGE. The eluting fractions containing the monomer peaks for WT and TM γS were next collected, concentrated as above for DLS and incubated for 6 days at 37 °C. Samples were then reanalyzed by SEC-MALS to detect aggregates that had formed from the monomers with time. The polydispersity index (PDI) was determined from the ratio of *Mw* to *Mn* (i.e. Mw/Mn), where *Mn* is the weight average number of molecules at each time interval. A monodisperse sample would have a PDI of 1.000. All values were calculated in ASTRA VII software and variability is a percentage of the average value under the protein peak (Wyatt Tech.). The percent of protein eluting in each peak was determined from the amount of protein recovered for that peak compared to the total amount of protein recovered in all the peaks using the in-line refractive index detector.

### Equilibrium chemical unfolding of WT and deamidated γS

Proteins were denatured with increasing amounts of guanidine hydrochloride (GuHCl) in buffer containing 100 mM NaPi (pH 7.0), 1.1 mM EDTA, and 5.5 mM DTT at 37°C overnight with a final protein concentration of 10 μg/mL. Samples were excited at 295 nm and emission spectra measured from 310-400 nm with slit widths set at 0.5 mm using a spectrofluoropolarimeter (Photon Technology Inc., NJ). The ratio of the 360 nm to 320 nm fluorescence intensities were used to calculate percent protein unfolded at each GuHCl concentration. Non-linear line fit was performed and data was best fitted to a sigmoidal two state unfolding curve using PRISM 6 (GraphPad Software, CA). The Gibbs free energy, ΔG_D_^0^ (kcal/mol), of unfolding was determined as previously described using the RTInK_D_ values calculated for K_D_=f_D_/f_N_ from the transition region of the GuHCl denaturation curves (28). The change in Gibbs free energy, ΔΔG_D_^0^ (kcal/mol), of unfolding was calculated using the difference between the ΔG_D_^0^ of WT (7.58 kcal/mol), and the ΔG_D_^0^ of each deamidated γS.

### Global hydrogen-deuterium (H/D) exchange of WT and deamidated γS

Proteins were incubated in 2.75 M GuHCl and pulse labeled for 1 min in D_2_O buffer. The number of amide hydrogens exchanged with deuterium was determined by measurement of the masses of whole proteins. Samples were diluted to 0.9 mg/mL in buffer containing 50 mM NaPi (pH 7.4), 300 mM NaCl, 10 mM imidazole, and 10 mM DTE. Diluted protein (10 μl) was mixed with 40 μl of 3.44 M GuHCl in H_2_O exchange buffer (20 mM NaPi (pH 7.4), 2.5 mM TCEP, and 1 mM EDTA). After unfolding for 0-32 min at 30 °C, 10 μL of each GuHCl-treated sample was mixed with 90 μL of D_2_O containing exchange buffer for 1 min at 30 °C. Labeling was quenched by addition of 25 μL ice-cold 0.8 M phosphate buffer (pH 2.4) and samples immediately frozen on dry ice. Masses of unexchanged proteins were determined by similar treatment, but by adding 90 μl of H_2_O exchange buffer instead. Mass analysis of γS was performed by individually and reproducibly thawing each sample and immediate injection onto an ice emerged 1 × 8 mm Opti-Trap™ protein trap cartridge (Optimized Technologies, Oregon City, OR) at 50 μl/min in a mobile phase containing 0.1% formic acid and 2% acetonitrile. After washing for 3 min, proteins were eluted in mobile phase containing 0.1% formic acid and 80% acetonitrile and mass measured using a LTQ Velos ion trap mass spectrometer (Thermo Fisher Sci.) using MS scans in enhanced mode. Spectra acquired during the approximately 30 sec protein elution were averaged, and measurements at each time point repeated 3 times. The region of the spectra from m/z 906-918 containing the +23 ion was used to calculate the number of deuterium exchanged into the various species at each time during GuHCl unfolding. The area of the 3 major peaks of deuterated protein were calculated and expressed as the percent of total area of all observed peaks.

Samples were also exchanged into D_2_O buffer as described above but without denaturant. The amount of deuterium uptake with time was measured by mass spectrometry using the methods above.

### Chemically-induced aggregation of WT and deamidated γS

Unfolding and aggregation induced by GuHCl were monitored using a BioTek Synergy 2 microplate reader (BioTek, Winooski, VT). Proteins were initially concentrated to 10 mg/mL using Ultra-15 Filter Units (10 kDa MWCO) and filtered with Millex-GV Syringe Filter Units (0.22 μm pore size). Samples were then prepared at 5 mg/mL in 25 mM NaPi (pH 7.2), 100 mM NaCl, with 10 mM DTT and 2 M GuHCl. Protein samples (60 μl) were transferred to black μCLEAR^®^ 96-well microplates (Greiner Bio-One, Monroe, NC; Catalogue No. 675096) which were then sealed with MiniStrips™ (Excel Scientific, Victorville, CA; Catalogue No. SP-2×8-50). Microplates were incubated with continuous shaking (1140 rpm) at 37 °C. Turbidity (405 nm), thioflavin T fluorescence (excitation/emission at 440±20/485±10 nm) and intrinsic fluorescence (excitation/emission at 284±5/360±20 nm) were measured every minute for 12 hours.

Aggregation kinetics were fitted to a single Boltzmann function using Origin 9.0 (OriginLab Corporation, Northampton, MA) and the fitting parameters were used to calculate the maximum rate of aggregation as previously described (29). Statistical significance was assessed using a two-sample t-test with unequal variances.

### Thermally-induced aggregation of WT and deamidated γS

Proteins were heated at 70 °C (i.e. the approximate midpoint of thermal unfolding(30)) and changes in turbidity measured. An aliquot of 100 uL of each crystallin was diluted to 24 uM using 100 mM NaPi (pH 7.4), 100 mM KCl, 1 mM EDTA, 2.5 mM TCEP and heated to 70 °C in Peltier Thermal Cycler-200 (MJ Research, CA). Turbidity of each solution was measured in 96-well plate as the change in optical density at 405 nm using Synergy-HT Microplate Reader (BioTek). Initial rates of aggregation were determined by linear regression using PRISM 6 (GraphPad Software) and p-values determined using t-tests.

For chaperone assays, either WT or TM were mixed with αA-crystallin (αA) in a ratio of 2:1, γS:αA, to a final volume of 100 μL containing a total of 100 μg of protein. Samples were diluted as needed with 100 mM NaPi (pH 7.4), 100 mM KCl, 1 mM EDTA, and 2.5 mM TCEP. Triplicates of each sample were prepared and incubated at 65 °C. Turbidity at 405 nm was measured on a plate reader at 0, 5, 10, 15, 30 and 60 minutes. Control samples contained 66.6 μg of WT or TM in 100 μL to match the amount of γS protein in experimental samples. To determine the identity of the precipitating species, samples were centrifuged, and the soluble proteins separated from the pellet. The pellet was washed, recentrifuged and the remaining insoluble proteins resolubilized. Both the soluble and insoluble proteins were analyzed by SDS-PAGE. After visualization with Coomassie blue staining, protein bands were excised and the protein identified by mass spectrometry.

## Results

### Surface deamidations increased protein aggregation of human γS under native conditions

Dynamic light scattering indicated that deamidation of γS increased its propensity to form large aggregates (Fig. 2). Freshly prepared WT γS at a concentration of 1 mg/mL, showed a predominant single peak with a mean *R*_h_ of 2.5 nm (1.5-4.9 nm range) and a smaller amount of scatter coming from a species between 10-100 nm in the intensity-weighted size distribution (Fig. 2 and Table 1). This is consistent with literature values of 2.0-2.5 nm for monomeric γS (30, 31). At a higher concentration of 5 mg/mL, similar *R*_h_ was detected for the main peak, but with a greater DLS intensity plus an additional species >100 nm (Fig. 2 and Table 1).

**Figure 2:**
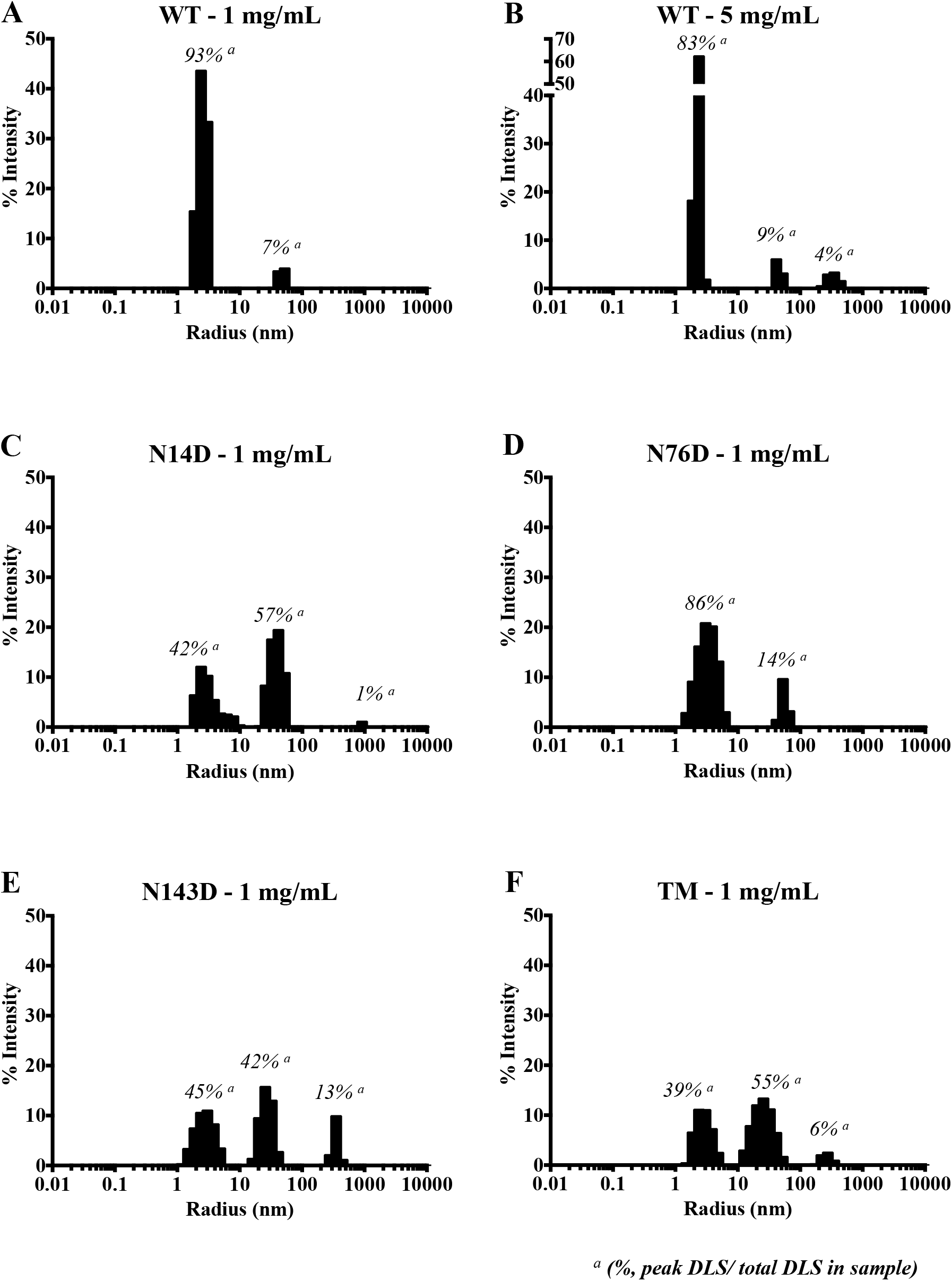
Effect of deamidation on radii of hydration, *R*_h_, of γS. The percent of dynamic light scattering intensity (DLS) was plotted for the size distributions between 1-10 nm (peak 1), 10-100 nm (peak 2), and 100-1000 nm (peak 3) of WT and deamidated γS. **A and B.** The % intensity of WT species at 1 mg/mL and 5 mg/mL respectively. **C, D, E and F.** The % intensity of N14D, N76D, N143D and N14D/N76D/N143D (Triple Mutant, TM) at 1 mg/mL. Data collected at 25 °C. ^***a***^The % DLS for each peak relative to the total DLS for that sample. Data are representative of three independent experiments.

**Table 1.** Sizes of γS in Peak 1 determined by dynamic light scattering

| γS | Concentration | Radius (nm) | % Pd | Mw-R |
| --- | --- | --- | --- | --- |
|  | mg/mL | Mean (SEM) | Mean (SEM) | Mean (SEM) |
| **WT** | 1 | 2.5 (0.2) | 19 (4) | 28.9 (3.26) |
|  | 5 | 2.4 (0.2) | 6* (2) | 25.0 (0.8) |
| **N14D** | 1 | 3.2 (0.5) | 37* (6) | 49.2** (5.6) |
| **N76D** | 1 | 3.3 (0.3) | 35* (3) | 55.0** (3.8) |
| **N143D** | 1 | 2.7 (0.2) | 29 (2) | 32.7 (3.6) |
| **TM** | 1 | 2.9 (0.3) | 22 (4) | 36.6 (2.5) |

In contrast to WT γS, the deamidation mimics showed predominantly two to three distinct peaks with a greater percent of intensity coming from the second and third peaks and less coming from the predicted monomer peak, peak 1. Peak 1 of the deamidated mimics had a *R*_h_ ranging from 2.7 to 3.3 nm (Table 1); while not significantly different from the WT, the N14D and N76D mimics’ Peak 1 had a wider size distribution, ranging from 1.5-9.9 nm and significantly greater polydispersity (%Pd) (35-37% compared to 19% for WT). Peak 2, with a *R*_h_ ranging from 10-100 nm, contributed more to the overall light scattering in the mimics than in the WT (14-57% compared to 7% for WT at the same concentration). In N143D and TM, peak 3 with a *R*_h_ between 200-400 nm was detected contributing 6-13% of the total light scattering.

The estimated *Mw*-R determined from unfractionated samples were greater for deamidated γS, particularly for N14D and N76D, than for the WT (Table 1). Since, the *Mw*-R is a weight average, even a small amount of a large species or nonglobular protein would increase the *Mw*-R more than the predicted *Mw* of 20.9 × 10^3^ g/mol. The wider distribution of *R*_h_, greater %Pd and estimated *Mw*-R of the deamidated mimics N14D and N76D suggested a small amount of a species larger than a monomer was present, even in Peak 1. All of the mimics had increased light scattering from peak 2 than WT, with N143D and the TM having an even larger species found in peak 3. However, the percent mass of these species predicted from reference globular proteins of similar *Mw* suggested their mass was less than 0.5%. Thus, a small number of large light scattering aggregates were detected in several DLS peaks in the deamidation mimics that were not present in WT γS.

### Surface deamidations increased Mw of human γS under native conditions

The formation of larger light scattering species by the deamidation mimics was confirmed by SEC-MALS on comparison of WT and TM γS (Fig. 3A). Both proteins eluted primarily as a monomer at 14.5 mL with a *Mw* of 20.5 × 10^3^ (± 0.2%) g/mol, comparable to the expected *Mw* of 20.9 × 10^3^ g/mol and with a polydispersity index of 1.000. In addition to the monomer peak, WT and TM γS exhibited an early eluting peak at 7.5 mL, overlapping with the void peak at 7 mL. This peak of TM was much higher in intensity and broader than that of WT. Similar results were obtained for other deamidation mimics (Fig. S1).

**Figure 3:**
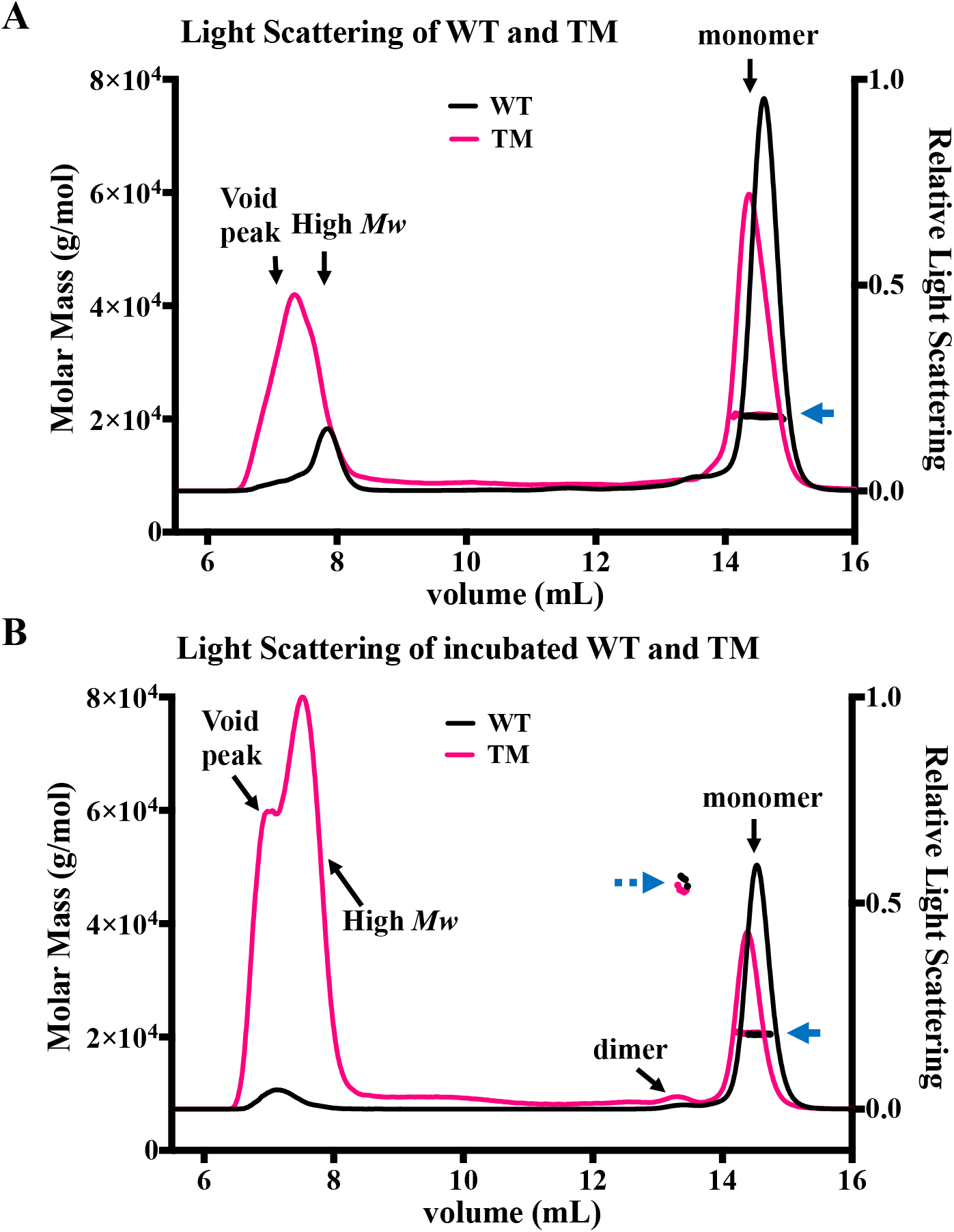
Relative light scattering (LS) of WT and deamidated γS. **A.** Light scattering of WT and TM γS during size exclusion chromatography in-line with multiangle light scattering (SEC-MALS). The right axis is relative Rayleigh LS of WT (solid, black line) and TM (solid, magenta line). The left axis is the molar mass (noted by blue arrows). Monomer peaks eluted at 14.5 mL and high *Mw* peaks eluted at 7.5 mL. Molar masses for the monomer peaks were 20.5 × 10^3^ g/mol (solid, blue arrow) compared to the predicted *Mw* of 20.9 × 10^3^. **B.** Light scattering of incubated WT and TM γS monomers. The monomer peaks isolated in panel A were incubated at 37 °C for 6 days and then reanalyzed by SEC-MALS. Monomer peaks were detected with 20.6 × 10^3^ g/mol (solid, blue arrow) and dimer peaks eluted at 13.5 mL with 46.8 × 10^3^ g/mol (dashed, blue arrow). Molar mass for the high *Mw* peak for the TM in panel B was estimated to range from 3-9 × 10^6^ g/mol (data not shown).

The monomer peaks of WT and TM γS shown in Fig. 3A were collected and their homogeneity verified by SDS-PAGE. The monomer peaks of each protein were then incubated at 37 °C for 6 days before re-examination by SEC-MALS (Fig. 3B). Incubating TM and, to a lesser degree, WT, generated a minor light scattering peak at 13.5 mL with a *Mw* of 46.8 × 10^3^ (± 1.2%) g/mol, corresponding to the approximate *Mw* of a γS dimer (Fig 3B). Moreover, the relative light scattering peak between 6 and 8 mL increased for both TM and WT, while the monomer peak decreased, suggesting the early eluting peak was a large oligomer of the monomer. This peak in TM has greater light intensity than the similarly treated WT γS.

The high *Mw* peak of TM had a *Mw* estimated to range from 3-9 × 10^6^ g/mol and contained less than 1% (0.8%) of the total protein. Even less protein eluted in this peak in WT preventing its accurate *Mw* determination. Both peaks were confirmed by mass spectrometry to contain γS. Hence, under native conditions the triple deamidation mimic more heavily favored a small population of conformer that was highly aggregation-prone.

### Effects of surface deamidations on the conformational stability of γS

Equilibrium unfolding experiments were performed to measure the thermodynamic stability of deamidated γS relative to WT γS (Fig. 4). There are two tryptophan (Trp) residues in the N-terminal domain (N-td) and two in the C-terminal domain (C-td) of γS. The location of these Trp residues permits their use as probes of the global structure (32). The Trps were selectively excited and their fluorescence measured in the presence of increasing concentrations of the denaturant, GuHCl. The midpoint of unfolding (C_M_) for WT was 2.58 M, comparable to the previously reported value of 2.3 M(33). The unfolding transition of WT was fitted with a two-state model and a Gibbs free energy of unfolding, ΔG°_N-U_, of 7.58 kcal/mol was derived. The C_m_ of the deamidated γS variants deviated only marginally from that of WT γS (Fig. 4). However, minor shifts in the early stages of the unfolding transition were observed for N76D and TM γS (Fig. 4), resulting in changes in ΔG°_N-U_ of −1.13 to −2.94 kcal/mol from that of WT. The trend towards a less stable protein at lower denaturant concentrations may reflect unfolding of the less stable N-td (33). The comparable Cm value with decreased ΔG°_N-U_ for N76D and TM indicates a decrease in the *m*-value (i.e. the dependence of ΔG°_N-U_ on denaturant concentration) which correlates strongly to changes in the amount of solvent accesible area exposed upon unfolding (34). Hysteresis was also observed upon refolding for all deamidation mimics and for WT γS (Fig. S2). The presence of hysteresis further suggested the tendency for the partially unfolded intermediates to aggregate with differing refolding trends between the proteins.

**Figure 4:**
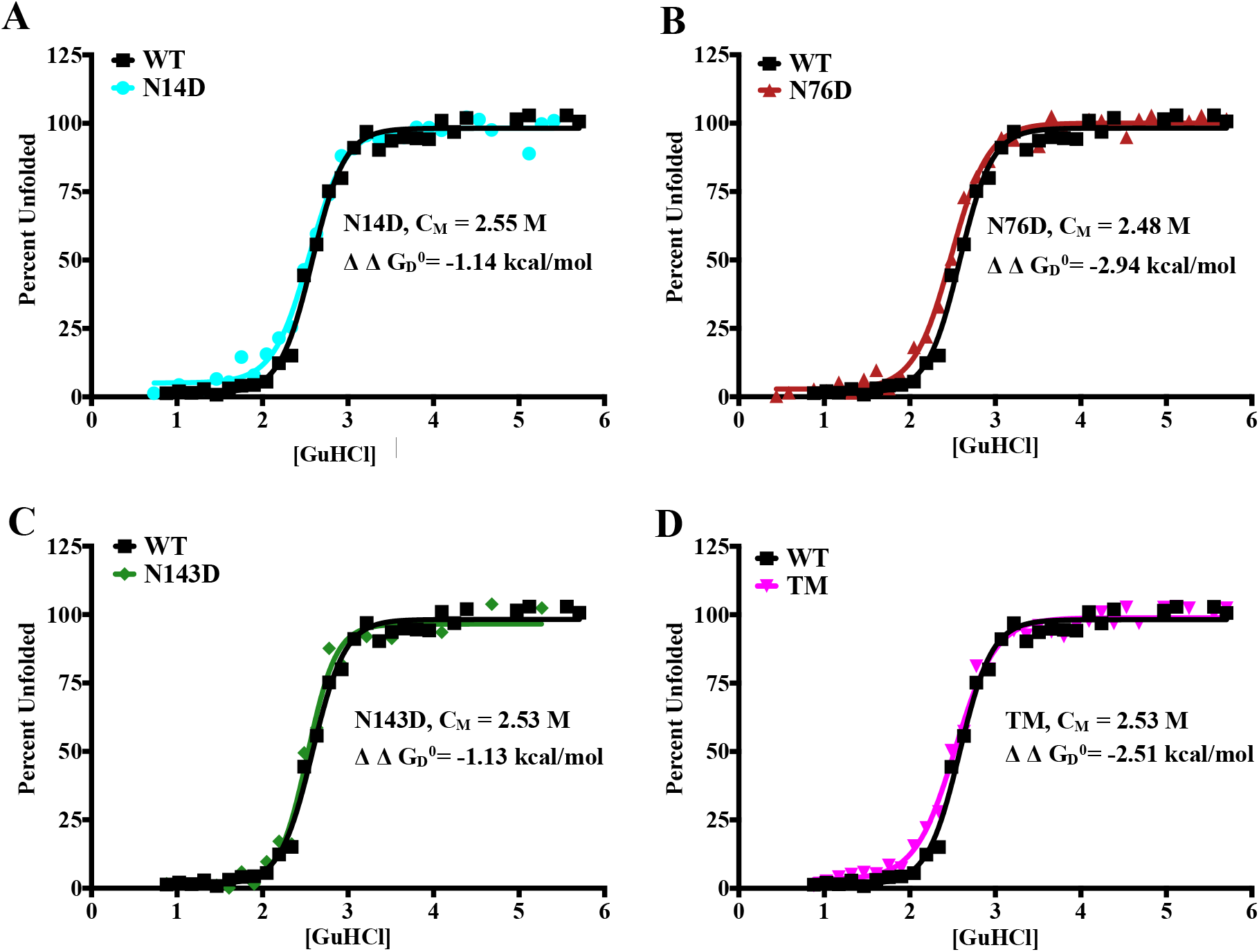
Equilibrium unfolding of WT and deamidated γS during incubation in GuHCl. Unfolding of γS was measured by changes in tryptophan fluorescence following an overnight incubation in increasing concentrations of GuHCl. **A-D.** Unfolding curves for N14D, N76D, N143D and TM, with the WT curve superimposed. The change in Gibbs free energy, ΔΔG_D_^0^ (kcal/mol), of unfolding was calculated using the difference between the ΔG_D_^0^ of WT (8.59 kcal/mol), and the ΔG_D_^0^ of each deamidated γS. The C_M_ for WT was 2.58 M.

### During chemical denaturation, partially unfolded intermediates of deamidated γS formed more rapidly than of WT γS

Proteins were unfolded in 2.75 M GuHCl, near the concentration of half equilibrium unfolding (Cm) determined in Fig. 4. Following different times of GuHCl exposure, they were subjected to pulse labeling in H/D exchange D_2_O buffer to measure the number of solvent exposed amides (Fig. 5). Three distinct major mass species were detected in WT following exposure to GuHCl for 16 min (Fig. 5A). This region of the spectrum from m/z = 906-918 contains the +23 charge state of the protein showing mass increases of ~43 Da (Peak 1), ~85 Da (Peak 2), and ~130 Da (Peak 3) in the GuHCl-treated protein, compared to the m/z of the unlabeled protein at 908.6 (arrows) (spectrum not shown). Peak 1, the major species observed immediately after addition of GuHCl (0 time), results from labeling the most solvent exposed amides in the native γS structure. In the spectra of TM (Fig. 5B), after 16 min of GuHCL exposure, Peak 1 was completely lost, and only Peaks 2 and 3 were observed. This indicated that TM unfolds more rapidly than WT, forming the partially unfolded intermediate in Peak 2, and the nearly fully unfolded form in Peak 3 at faster rates. Since the exchange was performed in 90% D_2_O, and back exchange was estimated to be approximately 10%, Peaks 1, 2, and 3, corresponded to the exchange of approximately 31, 62, and 94% of the protein’s backbone amides, respectively. The small peak to the right of the arrows, indicating the position of the +23 ion of the unlabeled proteins, may result from aggregated protein that is largely unavailable for H/D exchange.

**Figure 5:**
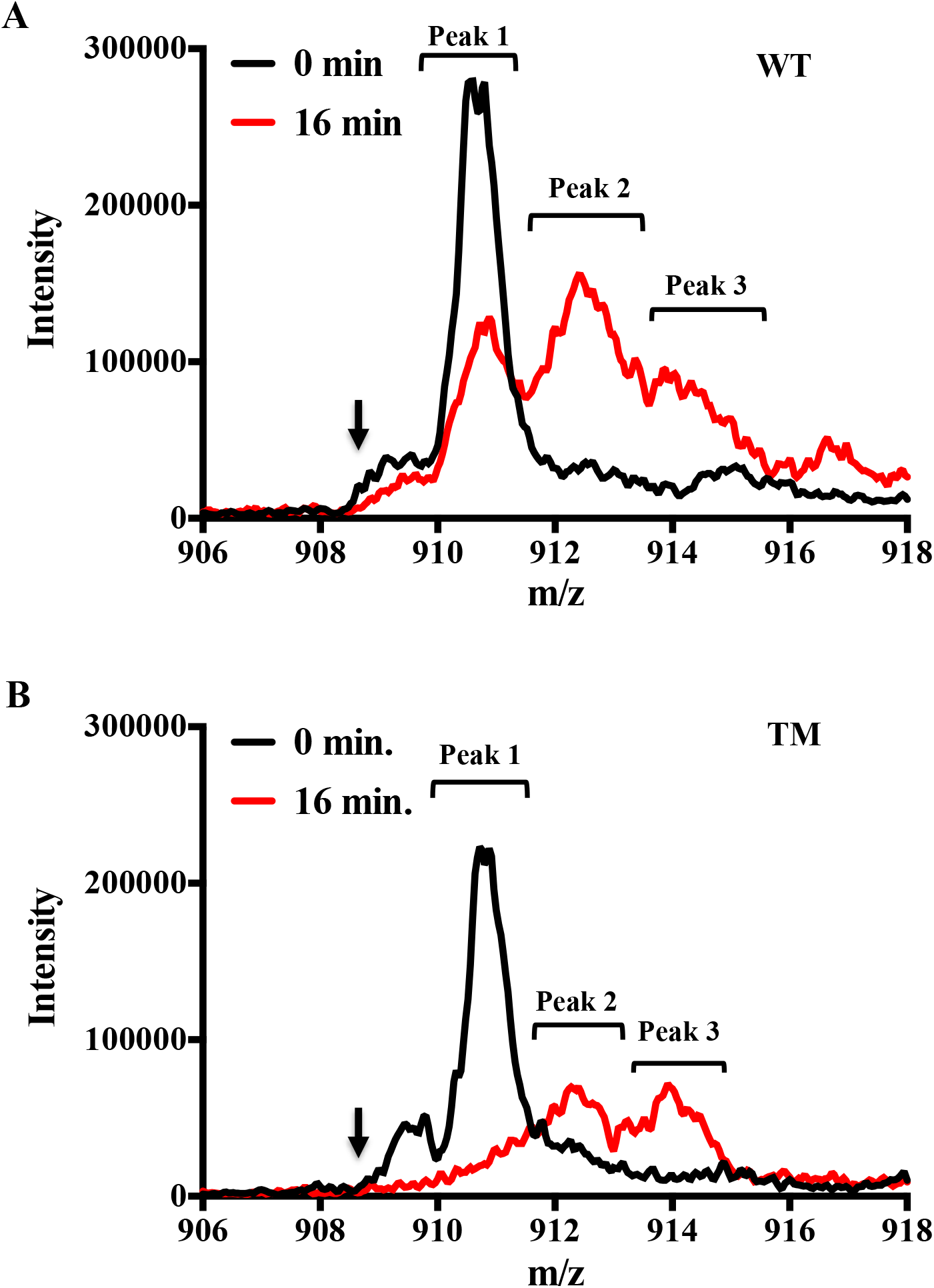
Deamidation increases the rate of γS chemical unfolding. Mass spectrum of the +23 charge state of WT and TM γS following partial unfolding in 2.75 M GuHCl. Proteins were unfolded in 2.75 M GuHCl for 0 and 16 min and subjected to pulse labeling in hydrogen deuterium (H/D) exchange D_2_0 buffer. Three major peaks were detected after 16 min in GuHCl with mass increases of 36-43 Da (peak 1), 83-85 Da (peak 2), and 120-130 Da (peak 3) over the unlabeled proteins (position of arrow, spectra not shown). **A**. The spectrum of WT at 0 min is overlaid with WT at 16 min. **B.** The spectrum of TM at 0 min is overlaid with TM at 16 min. In both panels A and B, there is a decrease in Peak 1 (folded species) and an increase in Peak 2 (partially unfolded intermediate) and Peak 3 (nearly fully unfolded species). Spectra are averaged from 3 independent experiments.

The percent of Peak 1 (native structure), Peak 2 (partially unfolded intermediate), and peak 3 (nearly fully unfolded protein) was calculated and the time course of their appearance and decrease after 0-32 min of exposure to 2.75 M GuHCl are shown in Fig. 6. At 4, 8, and 16 min, there was a significant decrease in Peak 1 (native structure) in N76D and TM proteins compared to the WT protein (Fig. 6A). The more rapid loss of native structure (Peak 1) in N76D and TM was accompanied by the significantly greater abundance of partially unfolded Peak 2 at 2, 4, and 8 min compared to WT protein (Fig. 6C), and a significantly lower abundance of Peak 2 at 32 min due to the faster accumulation of Peak 3 at 16 and 32 min (Fig. 6E). In comparison, N14D and N143D proteins unfolded at similar rates as the WT protein following GuHCl addition (Fig. 6B, D, and F). While, the identities of the exchanged amide residues in these three peaks were not determined, these results suggested that deamidation in N76D and TM γS led to more rapid production of partially and fully unfolded forms when γS was exposed to GuHCl. This observation was also consistent with their greater susceptibility to unfolding in GuHCL under equilibrium conditions (Fig. 4).

**Figure 6:**
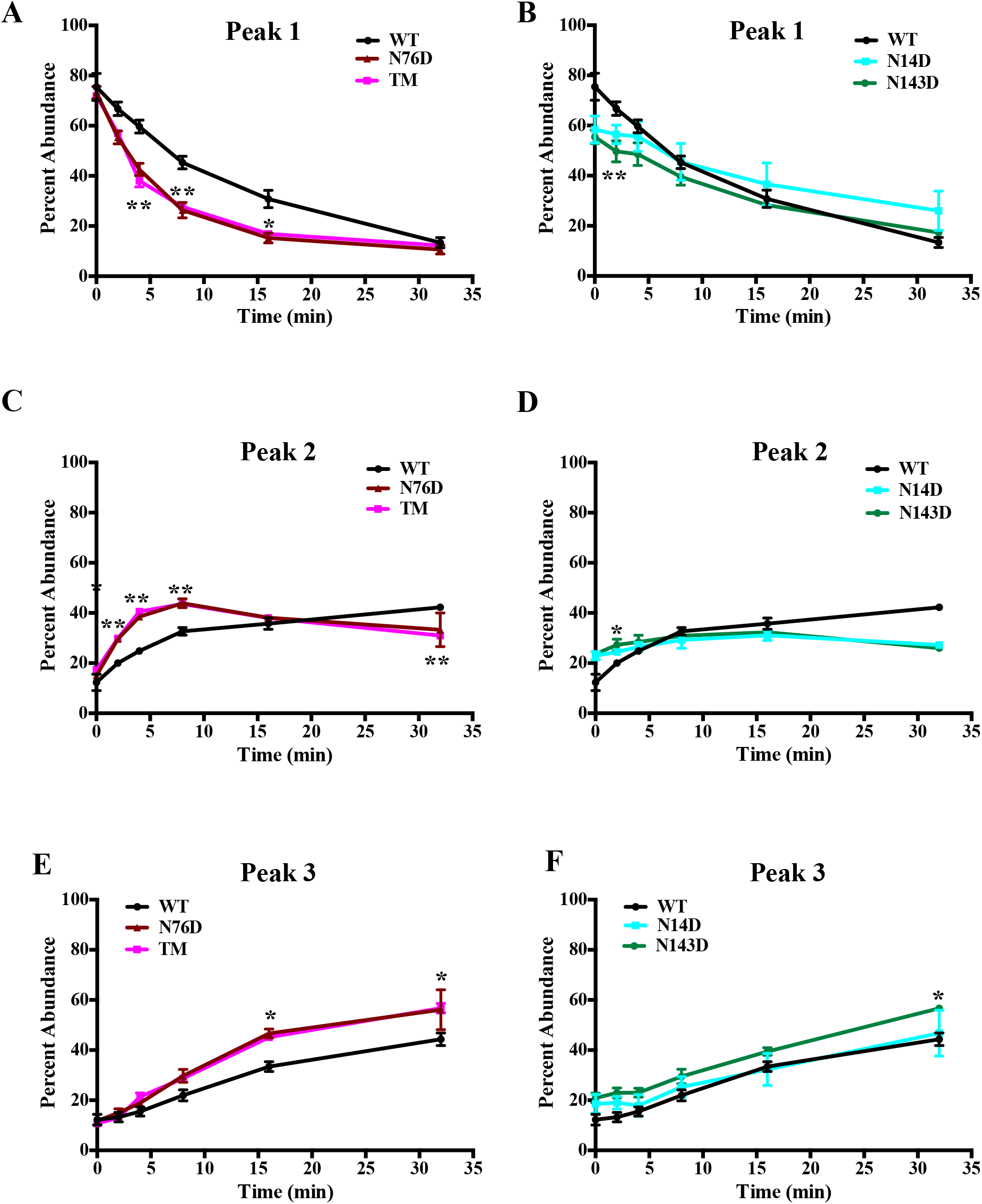
Changes in relative abundance of WT and deamidated γS intermediates during chemical unfolding. Percent area under the peak for each species of γS protein as shown in Fig. 5, calculated using raw spectra from pulse labeled protein in D_2_0 containing H/D exchange buffer. All data presented as percent abundance relative to total area of all peaks. Significant differences from WT with p-value < 0.01are denoted by ** and p-value ≤ 0.05 denoted by *, (n = 3).

### Surface deamidations increased chemically-induced aggregation of γS

To investigate whether deamidation of γS led to the formation of aggregation-prone, partially unfolded protein states, aggregation of WT and deamidated γS was monitored under conditions that facilitated unfolding. Proteins were incubated at 37 °C in the presence of 2 M GuHCl, i.e. a denaturant concentration corresponding to the beginning of the unfolding transition determined in Fig. 4. For WT there was a sigmoidal increase in solution turbidity and ThT binding over time which together indicated that the protein was forming aggregates with a degree of ordered fibrillar structure, as reported previously for γD-crystallin under comparable conditions (35). For N14D and N143D there was a minor but significant (p < 0.05) increase in aggregation propensity relative to WT, as shown by higher rates of aggregation (Fig. 7, inset) as well as higher levels of turbidity and ThT binding at the end of the incubation period (Fig. 7). In contrast, N76D showed a considerably greater aggregation propensity, consistent with its reduced stability (Fig. 4) and increased solvent accessibility under denaturing conditions (Fig. 6). The effects of deamidation at each site were accumulative with TM exhibiting the highest propensity to aggregate on the basis of aggregation rate. Despite the higher aggregation rate and final ThT binding, the solution turbidity of TM was disproportionally low, indicating a shift to more elongated (i.e. rod-like) structures (36). It is concluded that deamidation of γS, particularly at N76, promotes partial unfolding which thereby leads to enhanced aggregation.

### Surface deamidations increased thermally-induced aggregation of γS

The enhanced aggregation propensity of deamidated γS was further investigated under heat stress at 70°C (Fig. 8)(26). The N14D, N76D and TM mimics exhibited significantly greater turbidity after 5 minutes of exposure to elevated temperature when compared to WT and N143D γS (Fig. 8). Additionally, the rates of aggregation of N14D, N76D and TM were significantly greater than WT and N143D. The order of temperature stability from most to least stable was WT/N143D > N14D/N76D >TM. Deamidation at N14 and N76 located on the surface of the N-td affected the thermal-stability more than the deamidation at N143 on the surface of the C-td and the effects were cumulative, i.e. in TM.

**Figure 7:**
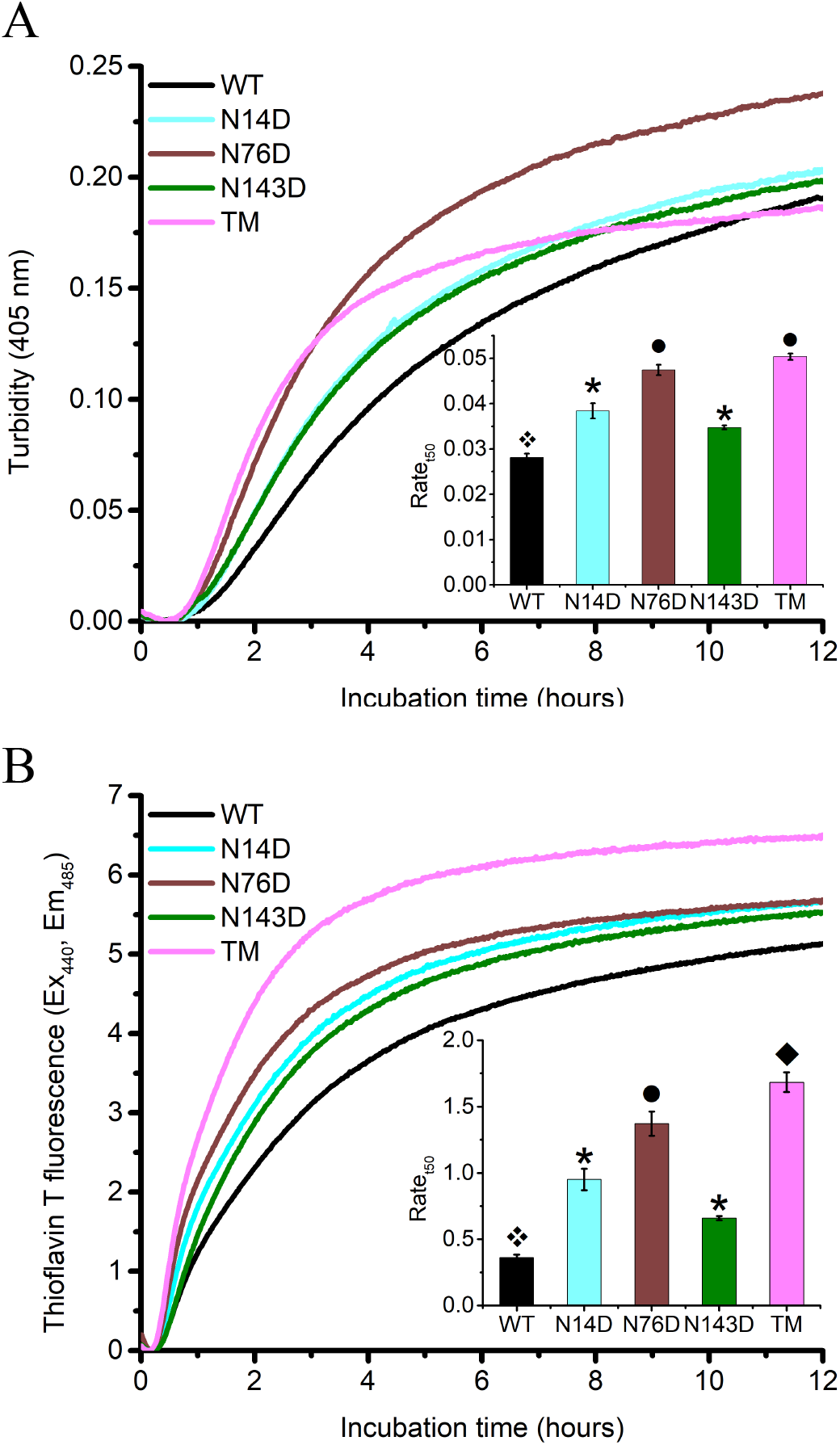
Effect of deamidation on chemically-induced unfolding and aggregation of γS. WT and deamidated γS at 5 mg/mL and in 2.2 M GuHCl were incubated in 96-microwell plates with shaking at 37 °C. **A.** Changes in turbidity at 405 nm. **B.** Changes in thioflavin T fluorescence at 440/485 nm were measured. Aggregation curves shown are average measurements (n ≥ 2) representative of three independent experiments. Inset shows maximum aggregation rate (Rate_t50_) derived from a Boltzmann sigmoid function (n ≥ 4, error bars = SEM). Rate_t50_ values that are significantly different from each other (p < 0.05) are distinguished by different symbols.

WT and TM γS were next incubated at 65 °C with and without the presence of a 2:1 molar subunit ratio of the molecular chaperone αA-crystallin (αA) to probe differences introduced from surface deamidations (Fig. 9). Both WT alone and WT in the presence of αA exhibited little change in turbidity with time as measured by light scattering at 405 nm over the course of the experiment. As expected, TM alone increased in turbidity over 15 minutes of heating, after which there was no further change. The lower turbidity value compared to Fig. 8 is due to the lower temperature that was used in that experiment. When incubated in the presence of αA, no increase in turbidity of TM occurred during the first 15 minutes, before aggregation commenced. Samples were analyzed by SDS-PAGE and mass spectrometry and it was determined that the precipitated proteins resulting from heating of TM alone and TM with αA, contained γS and γS plus αA, respectively (Fig. S4). The large error bar of the TM + αA sample at 60 min reflects the variability from the sample precipitating and large aggregates that formed. These results suggest that αA forms an unstable complex with deamidated γS that ultimately leads to rapid aggregation and co-precipitation.

**Figure 8:**
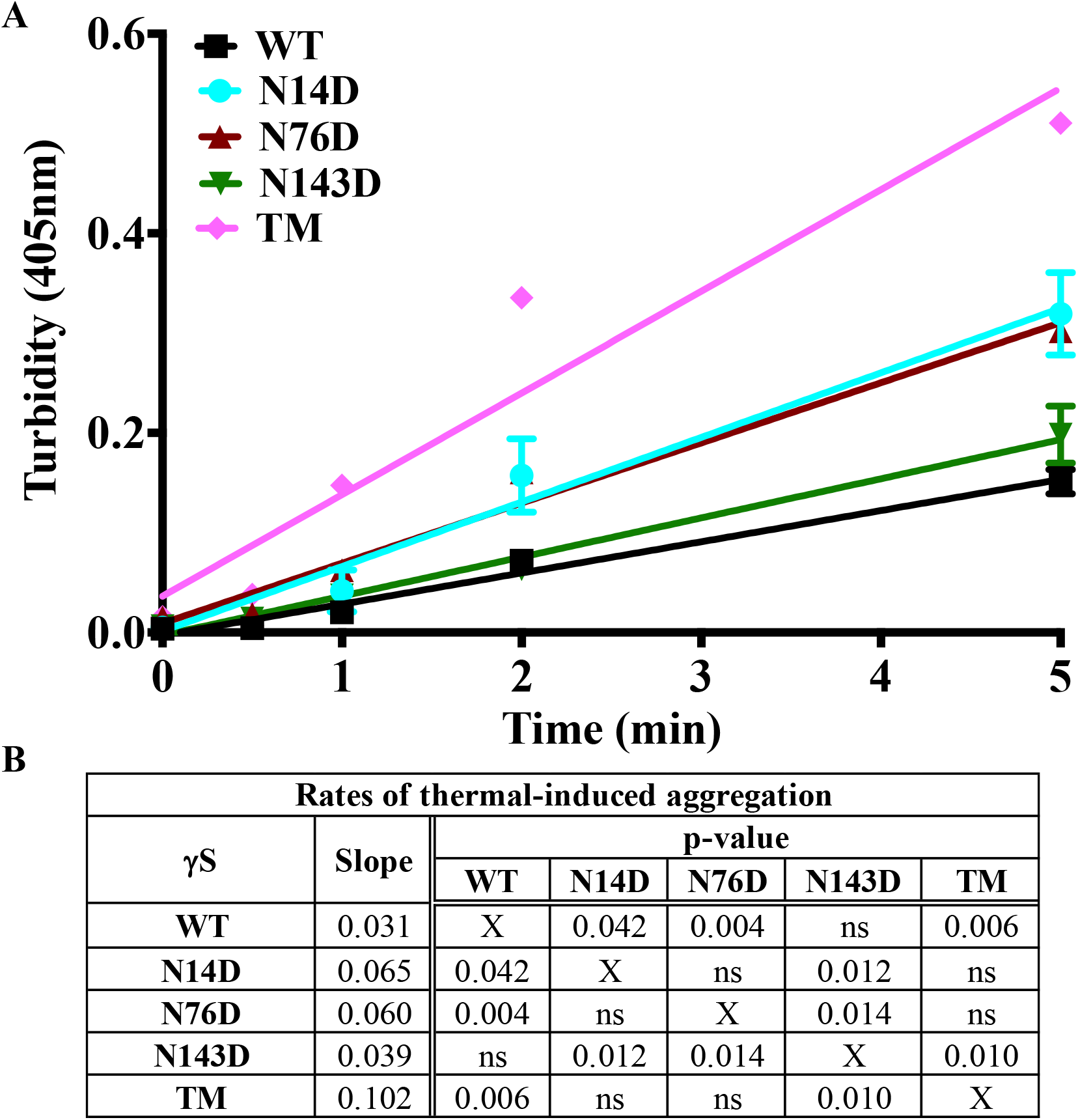
Deamidation increases thermal-induced aggregation of γS. **A.** Changes in turbidity during thermal-induced denaturation of WT, N14D, N76D, N143D and TM. Proteins at 24 μM concentrations were incubated at 70 °C and turbidity measured at 405 nm (n = 3, error bars = SEM). Note error bars are present on TM, but are not visible. Initial rates of aggregation were determined by linear regression. N14D, N76D and TM lines are significantly different than both WT and N143D. **B.** Table with p-values showing significant differences between aggregation rates for each γS (ns = not significant).

**Figure 9:**
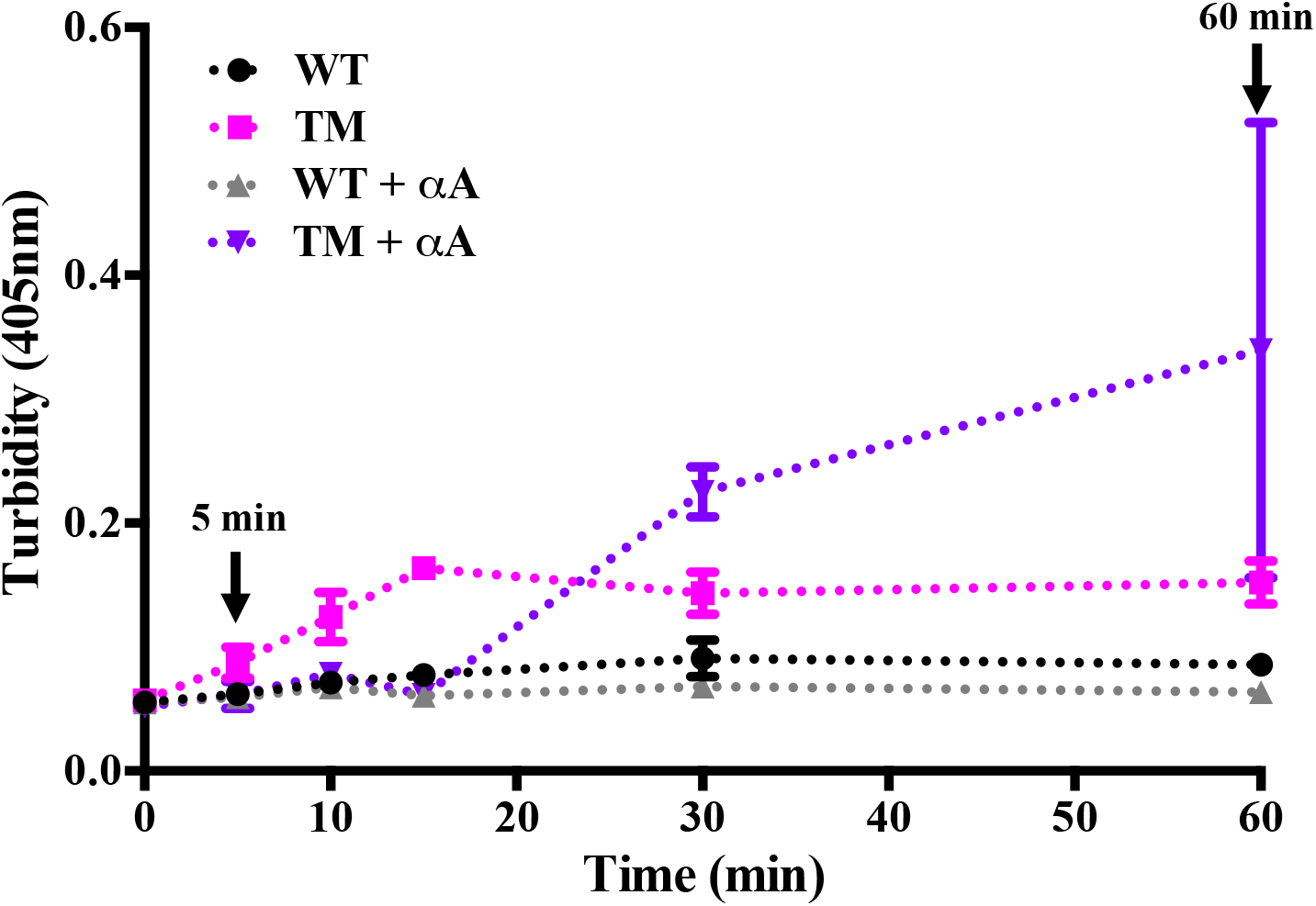
Thermal-induced aggregation of WT and TM γS with and without αA-crystallin. WT and TM with and without αA-crystallin were incubated at 65 °C and changes in turbidity at 405 nm were measured. (n = 3, error bars = SEM. Note: error bars are present on WT even though they are not visible at all time points and the large error bar on the TM at 60 min is due to sample precipitating.)

## Discussion

While cataracts are the major cause of blindness worldwide, the molecular mechanisms that lead to the formation of large light scattering aggregates that cause lens opacities are not known. The major proteins in the human eye lens, crystallins, are extensively modified during aging and cataract formation. Yet, a direct link between the accumulation of age-related modified crystallins and cataracts has not been well established. We present evidence here that one of the most prevalent modifications, deamidation of labile Asn residues, directly causes protein aggregation and present data that support a mechanism via the aggregation of partially unfolded intermediates as depicted in Fig. 10.

**Figure 10:**
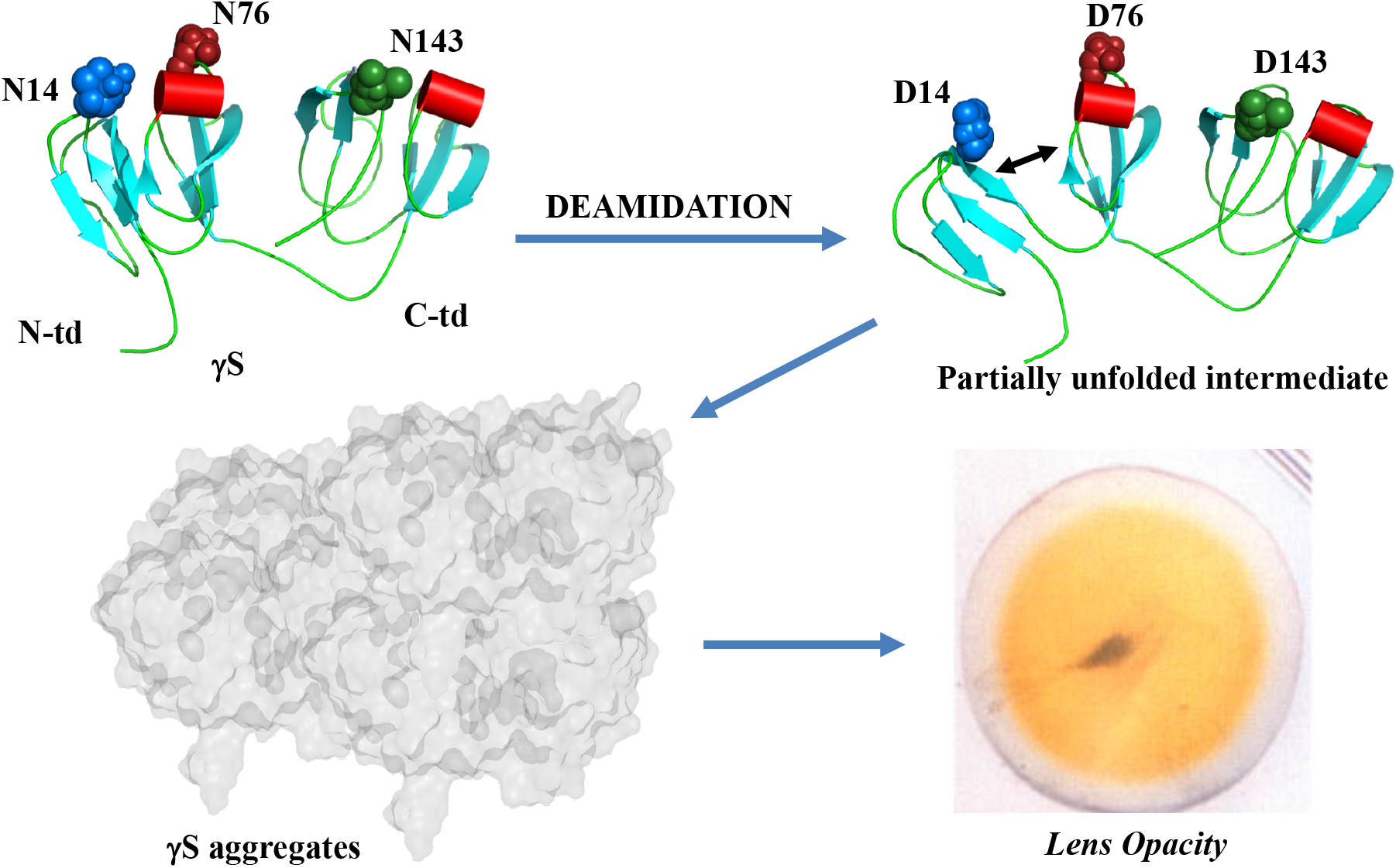
Model summarizing the effects of deamidation on γS. Deamidations associated with cataracts increase propensity of γS to partially unfold and form large light scattering aggregates that lead to lens opacity.

Deamidations at the labile Asn residues at positions 14, 76, and 143 in human γS led to large light scattering aggregates with a Rh greater than 10 nm that contributed 14-60% of the total scattering intensity compared to 7% for WT under the same conditions. Similar methods were used to detect light scattering in the P23T mutation in human γD-crystallin (similar to the P23V congenital cataract mutation), which showed < 5% of the total scattering intensity (37). Incubating the isolated γS monomer under physiological conditions led to increased aggregate formation with a corresponding decrease in monomer, supporting that deamidation directly led to the protein’s aggregation. It had previously been reported that deamidation at N76 and N143 together resulted in greater attractive interactions and that deamidation at N76 led to a tendency towards dimer formation without perturbing the structure or stability of γS (7). Taken together, these findings are consistent with deamidation leading to aggregation and thus, contributing to the insoluble deamidated crystallins isolated from cataractous lenses.

In these *in vitro* studies, the levels of deamidation-induced aggregation were < 1% of the total protein concentration, yet contributed to approximately half the light scattering intensity of the total protein depending on the deamidation site, i.e. only less than 1% of deamidated aggregate scattered a significant amount of light. In the crowded cellular milieu of the lens, this level of deamidation-induced aggregation is likely significant.

In order to determine if deamidation-induced aggregation was due to changes in stability, structures were probed during chemical and thermal-denaturation. Only small differences in equilibrium chemical-induced unfolding between WT and deamidated mimics were observed, suggesting that the overall structure of deamidated γS was comparable to that of WT (7, 26). Similar findings have been reported of none or little differences in the overall structure between several congenital cataract mutants (4, 5, 8, 9, 38–40). An emerging theme of the role of age-related modifications in crystallin structure is that these modifications do not appear to alter the overall structure of crystallins significantly *in vitro*.

However, during chemical kinetic unfolding of γS, a partially unfolded intermediate and a nearly fully unfolded intermediate were observed by H/D exchange that formed more rapidly in N76D and TM than WT. At a slightly lower GuHCL concentration the formation of these intermediates led to their rapid aggregation, suggesting that the partially unfolded intermediates were the species that were aggregating. The specific backbone amides undergoing rapid exchange in this intermediate have not yet been determined. The partially unfolded intermediate likely corresponds to the unfolding of the first Greek Key motif with approximately a 40 Da increase in mass in the N-td that was more prone to aggregate in the deamidated γS mimics. Others have reported that the N-td would unfold first followed by the C-td (33, 35, 41). Likewise, the N-td deamidations at N14 and N76 impacted γS more than the deamidation at N143 in the more stable C-td. Illustrating this is the observation that mutation at, at least, seven positions in γS is associated with early-onset cataract, whereas no inheritable mutations have so far been identified in the C-td (42).

During thermally-induced unfolding, deamidated crystallins rapidly aggregated supporting the chemical-unfolding findings. The mimic containing all three deamidation sites had the greatest effect on the structure of γS, with the N143D mimic in the C-td having only a minor effect on thermal-stability. Both deamidated mimics, N14D and N76D, in the N-td had a greater effect on thermal-stability than N143D in the C-td, further reflecting differences between the domains. Deamidated crystallins were also ineffectively prevented from precipitation by the chaperone action of α-crystallin with both proteins precipitating together (43–45). Similar effects of deamidation on the domain-domain interface in βB2 and γD have been reported (7, 21, 46). Our data suggest that once a threshold is reached where deamidation accumulates enough to unfold a protein, a cascade of nucleation-dependent aggregation and subsequent precipitation is initiated (47). More exposed hydrophobic regions would be present in deamidated γS than in WT for interaction with αA, given the lower temperature at which it aggregated. Several recent reports of deamidation at N76 in γS support these findings of decreased stability and ineffective chaperoning by α-crystallin due to the relative rapid aggregation of deamidated γS (43, 45, 48–50).

In earlier studies we detected changes in conformational dynamics in deamidated γS, which may help explain the results observed in the present study. These changes in dynamics were detected by differences in H/D exchange by NMR spectroscopy throughout the protein, i.e. at sites distant from the mutations including the β-helical regions in both the N- and C-tds (27). Noted changes were in β-strands in the first Greek Key motif near deamidation sites at N14 and N76. Increased exchange suggested increased solvent accessibility with a time frame of minutes to days of motion. More of the buried residues in these protein regions became exposed in the deamidated mimics during this time than in WT γS. The greater exposure of core residues and accessibility of other residues over time can explain the increased tendency to aggregate during both thermal and chemical unfolding of the deamidated mimics, particularly of the triple mimic.

γ-crystallins are stable, globular proteins that have considerable differences in Gibbs free energies between their native and unfolded states, mainly due to the energetically favorable burial of hydrophobic amino acid residues upon folding (33, 41). Complementary to this, the surface of γ-crystallins is dominated by hydrophilic and charged residues, particularly in loop regions, that form favorable hydrogen bonds with water, thereby providing solubility and stability (19). Human γS has ten Asp and fourteen Glu residues; two-thirds of these acidic residues have side-chain solvent exposures ranging between 40 to 80% with an average of ~55%, since solvation of charged residues is energetically favorable. Comparably, N14, N76 and N143 have solvent exposures of approximately 55, 58 and 60%, respectively. Ionized, aspartyl residues resulting from deamidation form stronger bonds with water than their uncharged, asparagine precursors. However, because surface residues are solvent-accessible in both native and unfolded forms of the protein, alterations in surface charge generally have a minor effect on net protein stability (51).

We attribute the increased aggregation propensity of the deamidation mimics to conformational fluctuations that expose otherwise slow-exchanging core residues to water, triggering unfolding and intermolecular association (27, 52). Additionally, and distinct from other sites of modification (e.g. N14, N143), deamidation at N76 may result in a non-native, surface salt bridge between the newly introduced anionic carboxylate and the cationic ammonium of R78. The minor increase in enthalpy accompanying salt bridge formation may come at a cost of conformational entropy as reflected by the conformational flexibility in the higher S^2^ values in the loop region intervening α-helix a2 and β-strand c2 and the reduced thermodynamic stability of N76D (Fig. 8) (27, 53, 54).

In summary, we have mimicked the aging process in crystallins by introducing deamidations via mutation. ‘Aged’ crystallins tend to partially unfold, exposing hydrophobic residues, as they accumulate deamidations. Exposed hydrophobicity leads to crystallin aggregation and the formation of large light scattering bodies that ultimately precipitate, causing the opacification of the eye lens associated with cataract (Fig. 10).

## Acknowledgements

The authors thank Wyatt Technology Inc. for use of the Dynapro Plate Reader.

## Funding Information

This research was supported in part by an Australian National Health and Medical Research Council Project Grant to J.A.C. (#1068087), by the National Institutes of Health Grants EY027012 to KJL and EY027768 to KJL and LLD. Mass spectrometry support includes Ophthalmology Proteomics Core NIH P30 EY010572 and S10 OD012246 to Oregon Health and Science University.

## Author Contribution Statement

CJV, DCT, CM, and KH performed experiments. LLD designed and performed mass spectrometry experiments and wrote methods. CJV, DCT and JAC designed experiments and wrote manuscript. KJL designed overall experiments and wrote manuscript.

## Abbreviations List

(TCEP): Tris(2-carboxyethyl)phosphine hydrochloride
(DTT): dithiothreitol
(SDS-PAGE): sodium dodecyl sulfate-polyacrylamide gel electrophoresis
(GuHCl): guanidine hydrochloride
(DLS, SLS, and MALS): Dynamic, static, and multi-angle light scattering
(H/D): hydrogen deuterium exchange
(MS): mass spectrometry
(WT): wild type
(TM): triple mutant
(N-td): N-terminal domain
(C-td): C-terminal domain

---

**Supplementary Figure 1:**
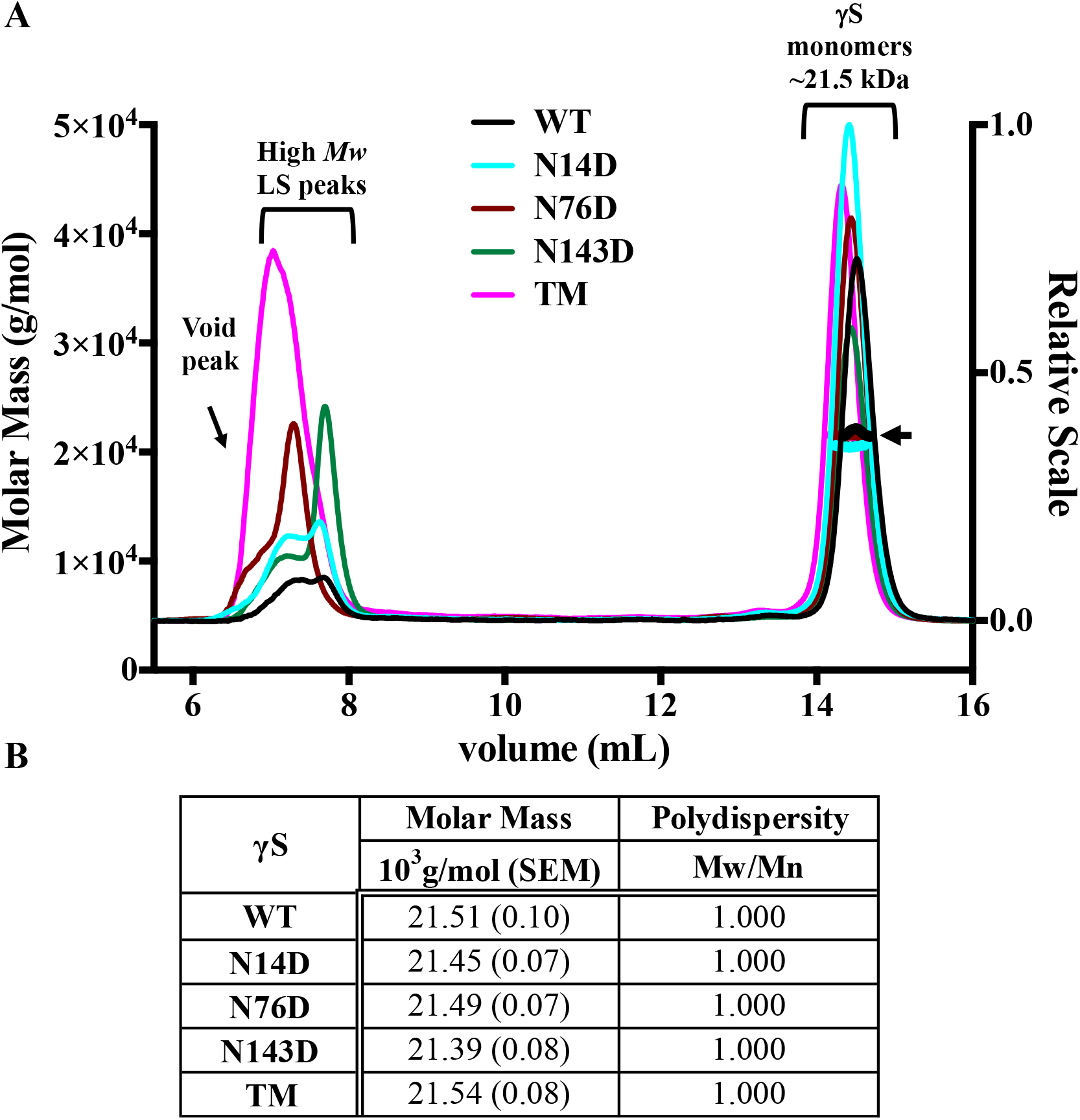
Increased relative light scattering of an aggregate γS species due to deamidation. **A.** A 50 μl sample of 5 mg/mL WT, N14D, N76D, N143D and TM were separated by size exclusion chromatography in-line with a multiangle light scattering detector and a refractive index detector (Superose 6 column, GE Health, IL in-line with a Dawn Heleos-II and Optilab rEX, Wyatt Technology, CA). Samples were prepared in 58 mM sodium phosphate (pH 6.8), 100 mM KCl, 1 mM EDTA and 1 mM DTT. Between 140 and 240 μg were recovered under the monomer peak eluting at 14.5 niL. The high molar mass peak eluted at 7.5 niL and was confirmed to contain γS with mass spectrometry. **B.** Molar masses and polydispersity of γS monomer from panel A. The molar mass of 21.5 × 10^3^ g/mol for all proteins is near the expected value of 20.9 × 10^3^ g/mol. The molar masses were not able to be determined for the aggregate, due to the low recovery of protein in this peak. Results were reproducible over multiple experiments.

**Supplementary Figure 2:**
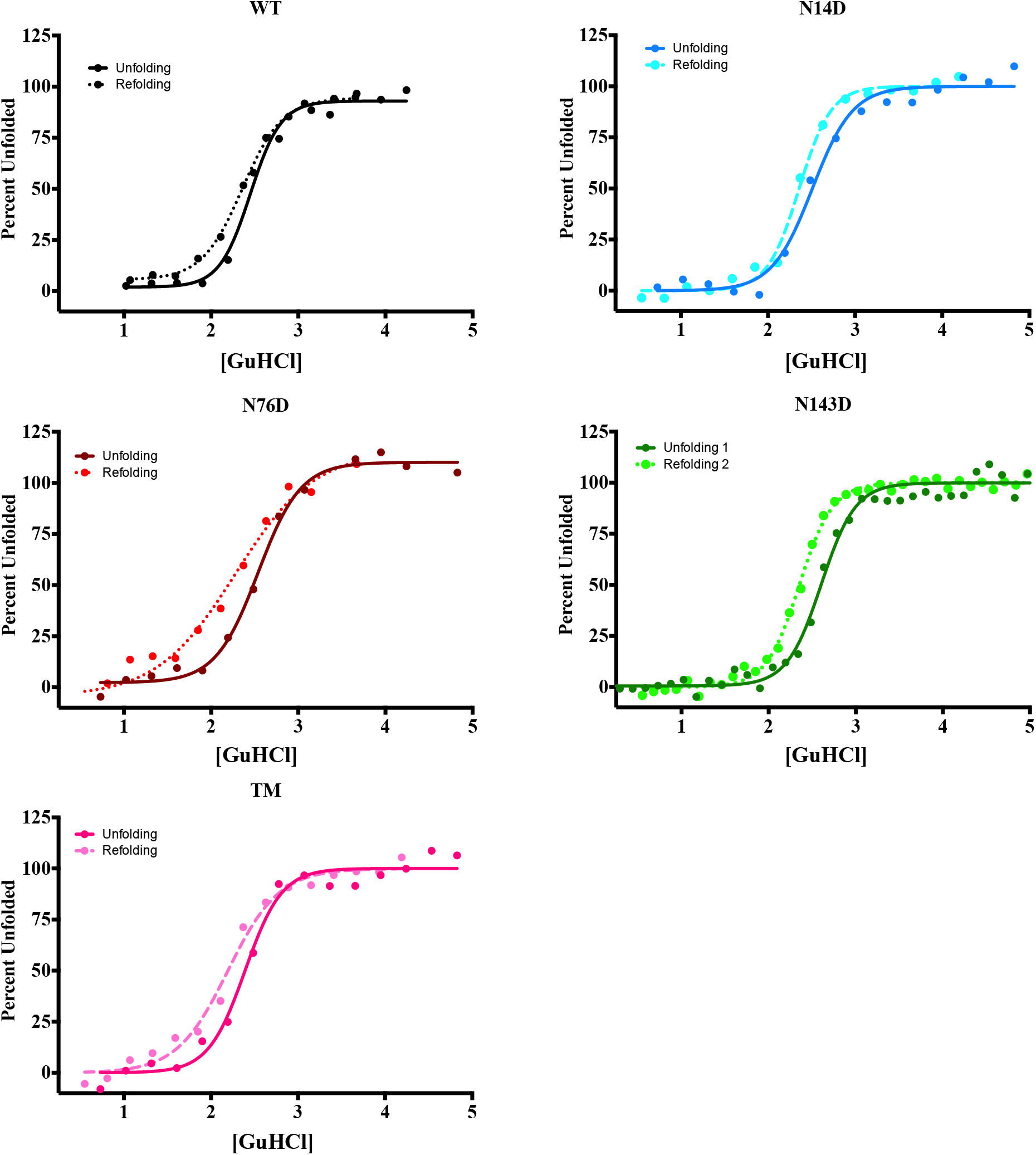
Unfolding/refolding of WT and deamidated γS showing hysteresis. Unfolding of deamidated mimics N14D, N76D, N143D, TM and WT γS crystallin were incubated in water bath at 10 ug/mL in 100 mM sodium phosphate, 1.1 mM EDTA, 5.5 mM DTT at 37°C for 12-16 hours with increasing amounts of guanidine hydrochloride (GuHCl). Refolding samples were unfolded as above, then diluted into decreasing concentrations of GuHCl and incubated at 37°C for 8-12 hours. Both sets of samples were then excited at 295 nm and emission spectra measured from 310-400 nm with slit widths set at 0.5 mm using fluorometer (Photon Technology Inc., NJ). The resulting 360 nm and 320 nm measurements were used to calculate percent protein unfolded at each GuHCl concentration and fitted to a sigmoidal two state unfolding curve using Prism 6 (GraphPad Software, CA).

**Supplementary Figure 3:**
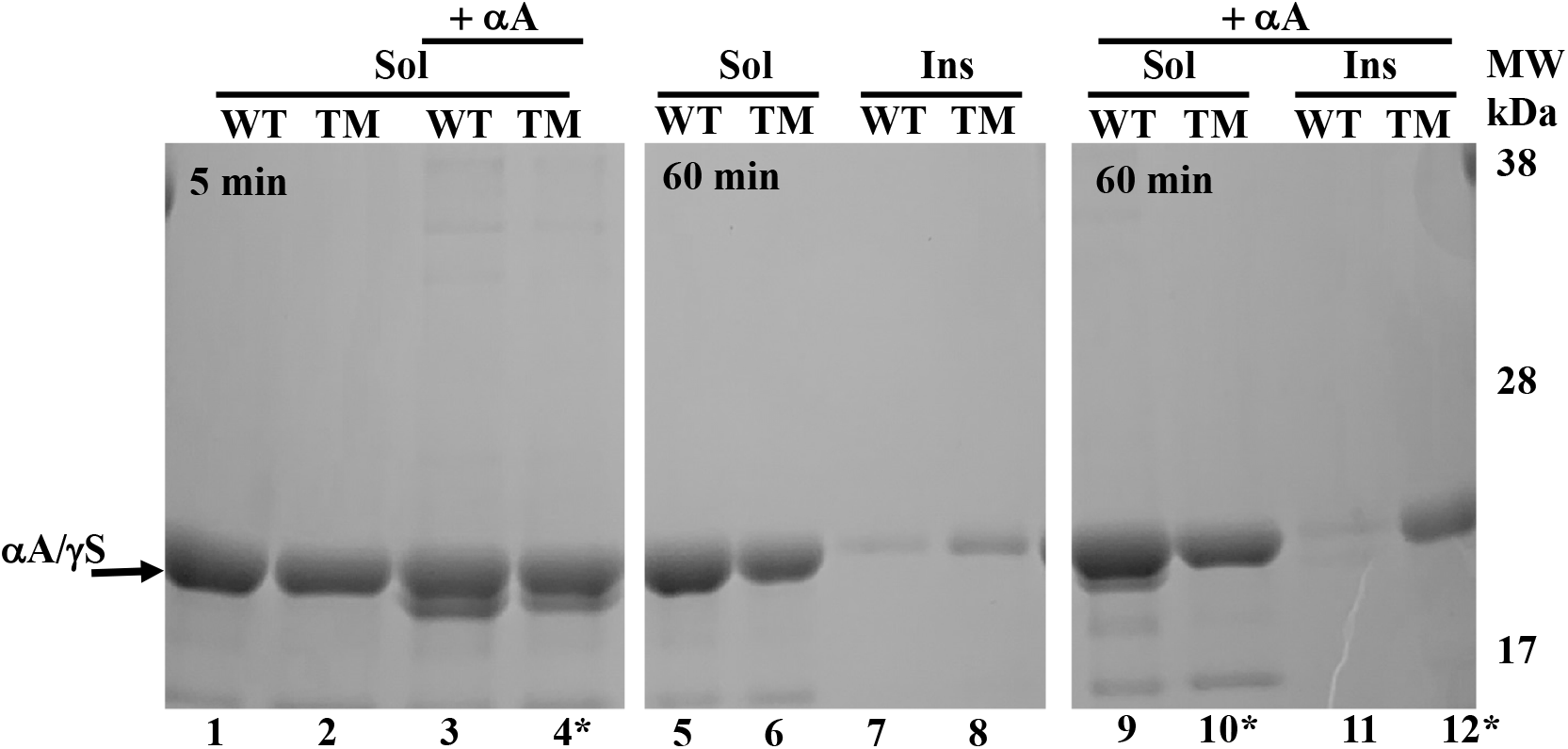
Thermal-induced aggregation of WT and deamidated γS with and without αA. SDS-PAGE of samples from 5 and 60 min of TM and WT incubated at 65 °C with and without αA chaperone in 100 uL containing 100 μg of protein with a ratio of 2:1, γS:αA. Lane 1-4, soluble protein after 5 min of heating; Lanes 5-8, Soluble (Sol) and insoluble (Ins) proteins after 60 min of heating; Lanes 9-12, Soluble and insoluble protein after 60 min of heating with αA. (*) Bands from lanes 4, 10, and 12 were confirmed by mass spectrometry to contain γS and αA, which co-migrate.

